# Blade-dependent molecular identity and neurogenic potential in the adult dentate gyrus

**DOI:** 10.64898/2026.07.24.740228

**Authors:** Unis Ahmad Bhat, Sergio Valbuena, Angel Marquez-Galera, Paula Tirado-Melendro, Marina Velotto, Carmen M. Navarrón, Frédéric Martins, Hélène Doat, Marlène Maitre, Vanessa Rouglan, Thierry Leste-Lasserre, Alexandre Brochard, Aixa V. Morales, Liset M. de la Prida, Djoher Nora Abrous, Jose P. López-Atalaya, Emilie Pacary

**Affiliations:** Univ. Bordeaux, INSERM, Neurocentre Magendie, U1215, F-3300 Bordeaux, France; Instituto de Neurociencias, Universidad Miguel Hernández-Consejo Superior de Investigaciones Científicas (UMH-CSIC), 03550, Sant Joan d’Alacant, Spain; Centro Neurociencias Cajal (CSIC), 28805 Alcalá de Henares, Spain

**Author notes:** These authors contributed equally (Sergio Valbuena, Angel Marquez-Galera). These authors contributed equally (Jose P. López-Atalaya, Emilie Pacary).

## Abstract

The dentate gyrus (DG) is a key hippocampal gateway for cognition and emotion and a major site of adult neurogenesis, yet its organization along the transverse (suprapyramidal– infrapyramidal) axis remains poorly understood. Here, by integrating bulk RNA sequencing of microdissected mouse DG blades with spatial transcriptomics and single-nucleus RNA sequencing, we define the suprapyramidal and infrapyramidal blades (SB and IB) as distinct molecular compartments characterized by anterior–posterior-dependent gene expression programs. Functionally, the SB exhibits enhanced neurogenic activity, particularly in the anterior DG, including increased progenitor proliferation and neuronal differentiation, whereas the IB contains a larger pool of quiescent neural stem cells. Together, these findings reveal molecular and functional specialization along both transverse and longitudinal axes of the DG and provide a framework for interrogating hippocampal subregional organization in health and disease.

## INTRODUCTION

The dentate gyrus (DG) is a key component of the hippocampal formation, serving as the main gateway for information entering the hippocampus. It receives inputs from the entorhinal cortex, processes this information, and relays it to the cornu ammonis (CA) areas^1^. This strategic position enables the DG to contribute critically to cognitive processing and affective regulation^2,3^. Moreover, the DG is one of the few brain regions where new neurons - dentate granule neurons (DGNs) - are continuously generated throughout life^4^, and alterations in DG neurogenesis have been associated with several mental disorders^5^, prompting extensive research into this unique region.

Traditionally, the DG has either been treated as a homogeneous structure or studied along the dorsoventral axis of the hippocampus. Along this axis, the dorsal hippocampus is primarily associated with learning and memory processes, whereas the ventral hippocampus is more strongly implicated in emotional and affective behaviors^3,6^, consistent with gradients in molecular, cellular, anatomical, and oscillatory properties^7^. In contrast, the transverse organization of the DG into suprapyramidal and infrapyramidal blades has been comparatively overlooked. The suprapyramidal blade (SB), also referred to as the upper or inner blade, lies above the CA3 pyramidal cell layer, whereas the infrapyramidal blade (IB), or lower or outer blade, is situated below the CA3 field^8^.

Nonetheless, growing evidence suggests structural and functional heterogeneity between these two blades. They differ in cellular composition and neural properties. Compared with the IB, the SB contains DGNs with greater dendritic length, higher spine density, distinct intrinsic electrophysiological features^9–13^, and a higher ratio of basket cells, suggesting differences in excitation–inhibition balance^14^.Their connectivity is also segregated: the lateral and medial entorhinal cortices preferentially innervate the SB and IB, respectively^15–17^, and the SB receives stronger supramammillary inputs^18^. In turn, IB mossy fibers preferentially target basal dendrites of CA3c neurons, whereas SB projections are biased toward apical dendrites in CA3a–b^19,20^. Functionally, activity-mapping approaches reveal consistent preferential SB activation during exposure to novel experiences^21–23^ and spatial learning tasks^24,25^. Despite these observations, the extent and functional significance of SB–IB differences remain poorly defined, underscoring a critical gap in our understanding of DG organization and function.

Here, we performed a comprehensive molecular characterization of the DG blades in adult mice. We precisely microdissected the two blades from both anterior and posterior regions of the brain (hereafter referred to as the longitudinal axis) and integrated bulk RNA-seq with spatial transcriptomics, single-nucleus RNA-seq, and functional analyses. We demonstrate that the DG is organized into molecularly distinct suprapyramidal and infrapyramidal compartments, with additional longitudinal specialization, associated with differential regulation of adult neurogenesis. Specifically, the SB displays higher neurogenic activity, whereas the IB is enriched in quiescent neural stem cells. Together, these findings establish a molecular framework for understanding how subregional specialization contributes to DG organization and function.

## MATERIALS AND METHODS

### Bulk RNA-seq

#### Animals

Five three-month-old *Ascl1^CreERT2^* ^+/-^;*Ai14* ^+/-^ male mice were used. *Ascl1Cre^ERT2^* (*Ascl1^tm1.1(Cre/ERT2)Jejo^*/J; RRID:IMSR_JAX:012882) and *Ai14* mice (B6.Cg- *Gt(ROSA)26Sor^tm14(CAG-tdTomato)Hze^*/J; RRID:IMSR_JAX:007914) were obtained from the Jackson Laboratory. Mice were maintained under standard conditions (12h light/dark cycle, light from 7:00 am to 7:00 pm, temperature 23 +/-1°C,) and received standard chow and water *ad libitum*. Mice were housed, bred, and treated according to the European directive 2010/63/EU and French laws on animal experimentation. All procedures involving animal experimentation and experimental protocols in this study were approved by the Animal Care Committee of Bordeaux (CEEA50) and the French Ministry of Higher Education, Research and Innovation (APAFIS authorization #38280).

#### Microdissection of DG blades and RNA extraction

Five brains were used for microdissection and subsequent RNA analysis. Coronal sections (50 µm) were cut from frozen brains in isopentane using a cryostat (CM3050 S Leica) at -20°C and mounted on polyethylene- naphthalate membrane 1mm glass slides (P.A.L.M. Microlaser Technologies AG) that were pretreated to inactivate RNases. Sections were then immediately fixed for 30 sec with 95% ethanol and incubated with 75% ethanol for 30 sec and with 50% ethanol for 30 sec. Sections were stained with 1% cresyl violet in 50% ethanol for 30 sec and dehydrated in 50%, 75% and 95% ethanol for 30 sec each, and finally 2 incubations in 100% ethanol for 30 sec were performed. Laser Pressure Catapulting (LPC) microdissection of the DG (subgranular zone and granule cell layer only) blades (Figure 1A) was performed using a PALM MicroBeam microdissection system v4.6 equipped with the P.A.L.M. RoboSoftware (P.A.L.M. Microlaser Technologies AG). Laser power and duration were adjusted to optimize capture efficiency. Microdissection was performed at 5X magnification. Microdissected tissues were collected in adhesives caps and resuspended in 250 µl guanidine isothiocyanate-containing buffer (BL buffer from ReliaPrep™ RNA Cell Miniprep System, Promega) with 10 µl 1-Thioglycerol and stored at −80°C until RNA extraction was done. For each brain, we began to microdissect the DG from the section at the anteroposterior coordinate -1.34 mm from the Bregma, when the two blades of the DG were visible. The SB blades of the first 25 sections (ranging from coordinates -1.34 mm to -2.57 mm relative to Bregma) were pooled to form the anterior suprapyramidal (AS) group, while the corresponding IB were pooled to create the anterior infrapyramidal (AI) group. Similarly, the SB and IB blades of the subsequent 25 sections (ranging from coordinates -2.57 mm to -3.8 mm relative to Bregma) were pooled to form the posterior suprapyramidal (PS) and posterior infrapyramidal (PI) groups, respectively.

**Figure 1:**
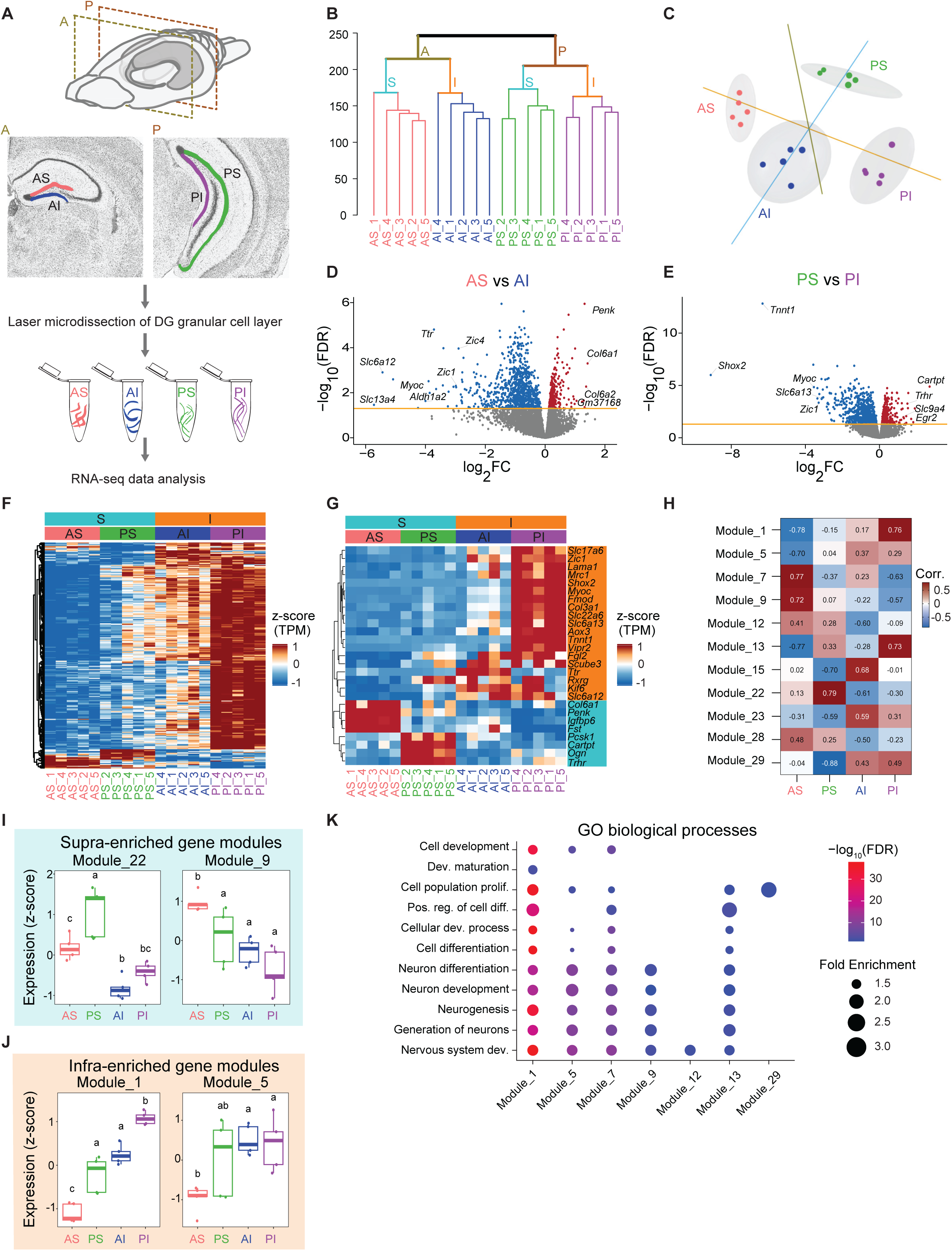
Transcriptomic profile of the DG along the transverse axis. **(A)** Experimental design. DG subregions (AS: anterior SB, AI: anterior IB, PS: posterior SB, PI: posterior IB) were microdissected and processed individually for bulk RNA-seq analysis (n=5 per subregion). **(B)** Hierarchical clustering of all samples based on TPM values after filtering lowly expressed genes. Samples segregate according to both anteroposterior position (A: anterior; P: posterior) and blade identity (S: supra; I: infra). **(C)** PCA based on TPM counts. Each dot represents one sample. Ellipses show 95% confidence intervals. **(D, E)** Volcano plots showing differential gene expression for AS vs AI (D) and PS vs PI (E) comparisons (FDR < 0.05). (F) Heatmap of consistently differentially expressed genes (|log_2_FC| > 0.5, FDR < 0.001) across blade comparisons (AS vs AI and PS vs PI). (G) Heatmap of candidate DG subregional marker genes based on TPM values. Genes enriched in IB are highlighted in orange (|log_2_FC| > 3), whereas genes enriched in SB are highlighted in turquoise (|log_2_FC| > 1). (H) Module–region association heatmap showing Pearson correlations between module eigengenes and DG subregions. **(I, J)** Box plots showing aggregated z-scores for modules enriched in SB (I) or IB (J). ANOVA determined significance; different letters (a, b, c, d) denote significant differences between groups (*p*-value < 0.05), groups with different letters are significantly different from each other. **(K)** GO biological process enrichment analysis for differentially expressed modules; dot color indicates FDR and size indicates fold enrichment.

Total RNAs were extracted from microdissected tissues using the ReliaPrep™ RNA Cell Miniprep System (Promega) according to the manufacturer’s protocol. The integrity of the RNA was checked by capillary electrophoresis using the RNA 6000 Pico Labchip kit and the Bioanalyzer 2100 (Agilent Technologies), and quantity was estimated using a DS-11 Spectrophotometer (DeNovix).

#### Bulk RNA-seq data analysis

Samples having RIN value > 8 were used for library preparation. Library for RNA sequencing was prepared using Illumina Stranded mRNA Prep, Ligation (Illumina) according to manufacturer’s protocol. In brief, mRNA was purified from 50 ng of total RNA using poly-T oligo beads. The purified mRNA was then fragmented, and cDNA was synthesized, followed by A-tailing and adapter ligation. PCR enrichment was performed for 14 cycles and sample was validated on the LabChip® GX Touch™ nucleic acid analyzer (Revvity). Libraries were sequenced on the NextSeq 2000 (Illumina) System using NextSeq 1000/2000 P2 Reagents (100 cycles) and paired-end reads (2 x 50 bases) according to manufacturer’s recommended protocols (∼ 20 million 50-bp paired-end reads per sample). The BaseCalls files obtained from the sequencer were converted into FASTQ files and demultiplexed using CASAVA_v1.8.2. Paired-end sequencing data with zero mismatches in the barcode sequences were considered, and raw data were checked for quality using FastQC v0.12.0^26^ and FastQ Screen v0.15.3^27^. The reads were mapped against the reference mouse transcriptome (GRCm39) using Hisat2 v2.2.1 with default parameters^28^. Only the reads that passed the quality filters (Phred quality score threshold of 30) were processed during the alignment. Genes were quantified for each sample using htseq-count from HTSeq package v0.11.1 with default parameters^29^. Genes/transcripts were filtered based on a complete match to the known transcripts. For exploratory analysis, raw counts were adjusted for batch effects using ComBat-seq v3.58.0^30^. Hierarchical clustering and principal component analysis (PCA) were carried out using log_2_TPM values of all detected genes (post library size normalization and low expressing transcript filtering). Complete-linkage clustering based on Manhattan distance was used to display sample relationships as a dendrogram. For differential gene expression analysis, library sizes were calculated for each sample and were used to normalize counts with the trimmed mean of M-values (TMM) method using calcNormFactors. Gene-wise dispersions were estimated and a generalized linear model was fitted using glmFit, with batch included as a covariate in the design matrix. Differential expression was assessed using the quasi-likelihood F-test (glmQLFTest) as described in edgeR package^31,32^. Statistical significance was determined by controlling the false discovery rate (FDR) using the Benjamini– Hochberg procedure and genes differentially expressed with a False Discovery Rate (FDR) value of < 0.05 were considered significant. R package ClusterProfiler v4.18.1^33^ was used to perform GO and other overrepresentation analysis. A Benjamini–Hochberg adjusted *p*-value cut-off of 0.05 was used to determine significant GO terms. For cell marker enrichment, the data set was downloaded from CellMarker 2.0^34^ and filtered for brain tissue before using for cell marker overrepresentation analysis in ClusterProfiler v4.18.1^33^.

#### Weighted gene co-expression networks

MultiWGCNA v1.9.1 package was used to construct the co-expression networks using the low expressing genes filtered counts per million data matrix (15,255 genes) of all samples^35^. The default network construction function constructNetworks was used to construct the coexpressing modules with the following parameters: networkType = “signed”, TOMType = “unsigned”, power = 9, minModuleSize = 30, maxBlockSize = 25000, minKMEtoStay = 0.7, mergeCutHeight = 0.20, deepSplit = 4. Differential module analysis among the DG subregions was carried out using the runDME function with ANOVA option to calculate p values on combined networks. For the identified differential gene modules, the relationship to DG subregion was analyzed with Pearson correlation, and GO analysis for biological processes was carried out using ShinyGO v0.85.1 web server with default parameters^36^.

### Quantitative real-time PCR

RNA was processed and analyzed following an adaptation of published methods^37^. cDNA was synthesized from total RNA by using QScript XLT cDNA Super Mix (Quanta Biosciences). qPCR was performed using a LightCycler® 480 Real-Time PCR System (Roche). qPCR reactions were done in duplicate for each sample, using gene-specific primers (Table 1), cDNA and LightCycler 480 SYBR Green I Master mix (Roche) in a final volume of 10 μl. PCR data were exported and analyzed in the GEASE software (Gene Expression Analysis Software Environment) developed in the Neurocentre Magendie (https://gease.neurocentre-magendie.fr/index.phtml). For the determination of the reference genes, the RefFinder method was used^38^. Relative expression analysis was corrected for PCR efficiency and normalized against two reference genes, the succinate dehydrogenase complex subunit (*Sdha*) and peptidylprolyl isomerase A (*Ppia*) genes. The relative level of expression was calculated using the comparative (2-ΔΔCT) method^39^.

**Table 1:**
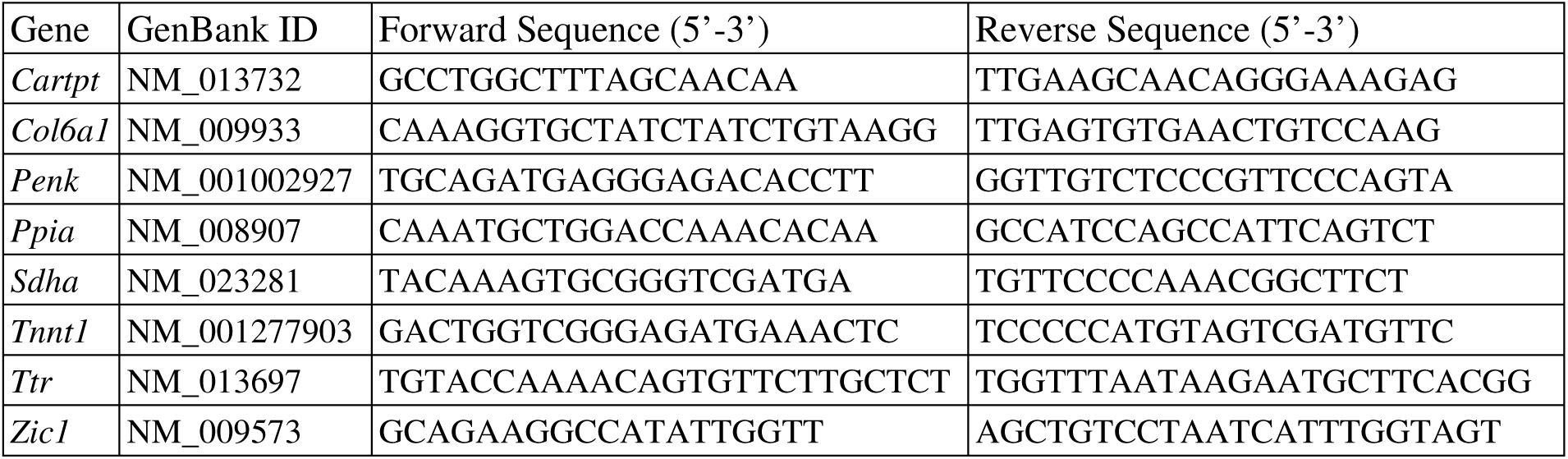
Sequences of primers used for qPCR.

### Spatial transcriptomics

#### Spatial transcriptomics dataset preparation

Slide-seqV2 spatial transcriptomic data^40^ from the DG granule cell layer (DG-sg) were obtained from Langlieb et al. (2023)^41^. Spots were annotated based on anteroposterior (AP) position (Anterior vs Posterior) and blade identity (Suprapyramidal (SB) vs Infrapyramidal (IB)). Prior to analysis, quality control filters were applied to remove spots with uncertain anatomical assignment. Specifically, spots corresponding to contralateral fragments or spatial outliers were excluded using spatial thresholds in Allen Common Coordinate Framework (CCF)^42^ coordinates (CCF_Z < 229.5 and CCF_Y > 200), to restrict analyses to anatomically reliable regions.

The AP classification was defined using a bregma reference of −2.57 mm, with pucks 39–52 assigned to the anterior region and pucks 53–65 to the posterior region. For blade annotation, SB regions were manually annotated for each puck based on DG-sg morphology. Spots were assigned according to blade shape, with a local bisecting line used only at the blade interface to separate SB and IB regions. After filtering and annotation, the DG-sg dataset comprised 29,858 spots, corresponding to 12,218 anterior and 17,640 posterior spots, as well as 12,599 IB and 17,259 SB spots. At a finer anatomical resolution, spots were distributed across compartments as follows: 5,571 Anterior–IB, 6,647 Anterior–SB, 7,028 Posterior–IB, and 10,612 Posterior– SB.

#### Spatial gene set scoring and contrast analysis

To project bulk RNA-seq–derived transcriptional signatures onto spatial transcriptomics data, gene set–based module scores were computed at spot resolution. For each contrast, the top 50 upregulated and top 50 downregulated genes (FDR < 0.05) were used to define directional gene signatures, where “up” corresponds to genes enriched in the reference condition (e.g. Anterior or SB), and “down” to genes enriched in the opposite condition (Posterior or IB), according to the direction of each contrast. Module scores were computed using Seurat’s v4.3.0.1 AddModuleScore^43,44^, yielding independent “up” and “down” scores per spot. Module score distributions (violin plots in figures) were compared using values >0 and two-sided unpaired Wilcoxon rank-sum tests, between anterior and posterior regions in A–P analyses and between suprapyramidal and infrapyramidal regions in S–I analyses. A composite contrast score was defined as: Contrast score = Score_up − Score_down. Spots from all pucks were projected into a pseudo-atlas by offsetting individual pucks along a common axis, to enable cross-puck visualization while preserving local spatial structure. Contrast scores were visualized using continuous color gradients with adaptive scaling. Statistical comparisons were performed using unpaired two-sided Wilcoxon rank-sum tests across relevant anatomical groupings. Additional metrics included the proportion of spots with positive or negative scores, as well as distributions of “up” and “down” module scores. Distributions were visualized using violin plots with embedded boxplots and jittered points.

### Single**-**nucleus RNA-seq (snRNA-seq)

#### Animals

To enable blade-specific dissection, we used *Calb1^Cre^*^+/-^;*Ai9*^+/-^ mice, in which DGNs are robustly labeled with tdTomato following trimethoprim (TMP) administration^45^. Two adult (five-month-old) male mice were used for snRNA-seq. These mice were generated by crossing *Calb1-2A-dgCre-D* mice (RRID:IMSR_JAX:023531) with the *Ai9* reporter line (RRID:IMSR_JAX:007905). To stabilize dgCre recombinase and permanently label DGNs with tdTomato, mice received a single intraperitoneal injection of TMP (100 mg/kg) at 1 month of age. Animals were housed in ventilated cages at a standard pathogen free (SPF) facility with *ad libitum* access to food and water, maintained at controlled temperature (23 °C) and humidity (40-60%) under 12h light/dark cycle. Experiments and procedures were conducted in accordance with the European directive 2010/63/EU and Spanish laws for animal research, and were approved by the CSIC ethics committee and authorized by the Comunidad de Madrid (PROEX162-19) and Generalitat Valenciana (2024-VSC-PEA-0001).

#### DG dissection and nuclei isolation

Animals were sacrificed by cervical dislocation, and brains were rapidly extracted. Coronal brain sections (300 µm thick) were obtained using a vibratome in ice-cold 1× HBSS. To minimize potential circadian effects, dissections were performed before noon.

For each brain, four coronal sections containing the anterior DG were manually dissected to separately isolate SB and IB. Dissected tissue was collected in 400 μl of ice-cold Magnetic-Activated Cell Sorting (MACS) buffer (0.5% BSA, 2 mM EDTA in PBS 1x). DG sections were transferred to a dounce tissue grinder containing 400 μl of MACS buffer and mechanically homogenized 12-15 times. The resulting suspension was transferred to 2 ml Eppendorf tubes and centrifuged at 500 G for 15 min at 4°C. The pellet, containing the DG cells, was resuspended in 500 μl of ice-cold lysis buffer (10 mM Tris-HCl, 10 mM NaCl, 3 mM MgCl2 and 0.1% IGEPAL) and kept in ice for 5 min to release nuclei. Samples were centrifuged again, at 500 G for 30 min at 4°C. The nuclei pellet was resuspended in 1% BSA in PBS 1x. For each sample, 15,000 nuclei were purified by flow cytometry using a BD Biosciences FACS Aria III.

#### Library preparation and sequencing

Purified nuclei from SB and IB of the DG were used to generate cDNA libraries. Briefly, nuclei were loaded into a Chromium Single Cell A Chip (10x Genomics) and processed according to the manufacturer’s instructions. snRNA-seq libraries were generated using the Chromium Single Cell 3’ Library & Gel Bead Kit v2 together with the i7 Multiplex kit (10x Genomics). Final libraries were pooled and sequenced on an Illumina HiSeq 2500 sequencer to obtain 75bp paired-end reads.

#### snRNA-seq data analysis

snRNA-seq data were processed using a standardized computational workflow. Raw sequencing reads were subjected to quality assessment with FastQC (Babraham Institute) and MultiQC (Seqera). Alignment, barcode processing, and UMI counting were performed using Cell Ranger v7.2.0 (10x Genomics). Ambient RNA contamination and spurious counts were subsequently corrected using CellBender v0.2.0 (Broad Institute). A total of 8,414 nuclei were recovered, with a mean sequencing depth of 69,505 reads per nucleus and an average of 809 detected genes per nucleus.

Downstream analyses were conducted in R using Seurat v5.1.0^46^. Nuclei with >1% mitochondrial gene content were excluded to ensure high-quality nuclear transcriptomes. Data normalization was performed using the global-scaling LogNormalize method with a scale factor of 10,000. SB and IB datasets were merged using the Merge function in Seurat. Highly variable genes were identified with FindVariableFeatures, and normalized expression values were scaled using ScaleData with default parameters. PCA was conducted on the top 2,000 most variable genes. Inspection of the leading principal components did not reveal evidence of batch effects.

Graph-based clustering was performed by constructing a shared nearest neighbor (SNN) graph using FindNeighbors based on the first 10 principal components, followed by community detection with FindClusters at a resolution of 0.4. This analysis identified 15 clusters. Two small clusters corresponding to oligodendrocytes and astrocytes were manually merged with their respective major populations. A minor population of ependymal cells detected in one infrapyramidal sample was excluded from further analysis. Four clusters were annotated as DGNs based on enrichment of canonical markers, including *Prox1* and *Stxbp6*. The remaining clusters comprised major glial populations, including astrocytes, oligodendrocytes, and microglia.

Cluster-specific marker genes were identified using the FindAllMarkers function in Seurat. Low-dimensional embedding and visualization were performed using Uniform Manifold Approximation and Projection (UMAP)^47^ computed on the first 10 principal components. Differential expression analysis (DEA) between cell populations was conducted using the FindMarkers function, comparing groups based on cluster identity, anatomical blade (SB vs. IB), or their interaction. P-values were adjusted for multiple testing using the Benjamini– Hochberg false discovery rate (FDR) correction, and significance was determined according to adjusted FDR thresholds.

### Bromodeoxyuridine and tamoxifen injections

Three-month-old *Ascl1^CreERT2^*^+/-^;*Ai14*^+/-^ mice were injected with a single dose of bromodeoxyuridine (BrdU, 10 mg/ml, i.p., 100 mg/kg body weight) and sacrifice 2h later (proliferation) or 4 weeks later (neuronal differentiation).

To label mature adult-born DGNs (dendrite analysis), Three-month-old *Ascl1^CreERT2^* ^+/-^;*Ai14* ^+/-^ mice were injected with a single dose of tamoxifen (120mg/kg, i.p.) and sacrifice 12 weeks post-injection.

### Tissue processing

Mice were deeply anesthetized with an intraperitoneal injection of a mixture of pentobarbital (300 mg/kg; Exagon®, Axience) and lidocaine (30 mg/kg, Lidor®, Axience) before transcardial perfusion with ice-cold phosphate buffer saline (PBS, 0.1M, pH=7.3) and then with ice-cold 4% paraformaldehyde (PFA) in PBS. Brains were post-fixed 24h in 4% PFA before being cut coronally and serially (10 series of 40 µm sections or 3 series of 100 µm sections for dendrite morphological analysis) with a vibratome (Leica). One series was used for each analysis.

### Immunohistochemistry and analysis

#### Chromogenic immunohistochemistry for proliferation and dendrite length analyses

Endogenous peroxidase activity was first quenched by incubating the sections in 1.2% hydrogen peroxide in methanol for 30 min. For BrdU detection, an additional step was carried out with a 30 min incubation with 2N Hydrochloric acid at 37°C. Prior to primary antibody incubation, sections were incubated with blocking solution (3% normal serum and 0.3% Triton X-100 in PBS) for 45 min at room temperature. Sections were incubated overnight with the primary antibody (mouse anti-BrdU, Santa Cruz Biotechnology, sc-32323, 1/100; rabbit anti-Tomato, Ozyme, 632496, 1/500) in blocking solution followed by an incubation with a biotin-labeled secondary antibody for 1.5 hour on the following day. Biotin-streptavidin technique (ABC kit; Vector Laboratories; PK-4000) with 3, 3′-diaminobenzidine (DAB) as chromogen was used to label the samples permanently for visualization of immunoreactivities.

BrdU-positive cells were quantified in six 40 µm sections from a single series spanning the SB and IB, with the first three sections corresponding to the anterior DG and the last three to the posterior DG. For dendrite length measurements, dendrites were traced using a 100× 1.4NA objective and a semiautomatic neuron tracing system (Neuron Tracing from Neurolucida^®^ software; MicroBrigthField) as previously described^48^.

#### Fluorescent immunohistochemistry for neuronal differentiation and NSC analyses

For neuronal differentiation analysis, a step of antigen retrieval was first performed by incubating sections in 10 mM sodium citrate (pH 6) for 2 min at 100°C, followed by incubation in 2 N HCl for 30 min at 37°C. Sections were blocked (5% normal serum, 0.3% Triton X-100 in PBS) for 1h and incubated overnight with primary antibodies (mouse anti-BrdU, Millipore NA61, 1:400; chicken anti-NeuN, Millipore ABN91, 1:1000). For NSC analyses, sections were permeabilized (1% Triton X-100 in 0.1 M PB) and blocked (10% goat serum, 0.25% Triton X-100 in 0.1 M PB), followed by antigen retrieval in 10 mM sodium citrate at 80°C for 20 min and overnight incubation with primary antibodies: rabbit anti-GFAP (Dako, Z0334, 1/1000), mouse anti- MCM2 (Bd Biosciences, 610701, 1/1000) and rat anti-Sox2 (Invitrogen, 14-9811-82, 1/400). After washes, sections were incubated with Alexa Fluor–conjugated secondary antibodies (1:1000), counterstained with DAPI (1:10,000), and mounted with Aqua Polymount.

Images were acquired on a Leica SP5 or SP8 confocal microscope (z-step: 2 µm). Six DG sections per animal were analyzed as described previously. Quantifications were performed manually and blind using ImageJ/Fiji. NSCs were identified by Sox2/GFAP expression and radial morphology in the subgranular zone (SGZ), with proliferative NSCs defined by MCM2 expression. Granule cell layer (GCL) area was measured using DAPI staining.

### Statistics

GraphPad Prism was used for statistical analyses. No statistical methods were used to predetermine sample sizes; however, our sample sizes are comparable to those generally employed in the field. Results are presented as mean ± s.e.m (standard error of the mean) in Figures 3C-E and 4G and a two-tailed paired Student’s t-test was used in these figures. Statistical tests and sample size (n) are indicated in the figure legends. Statistical values are mentioned in the text, legends, figures or tables.

**Figure 2:**
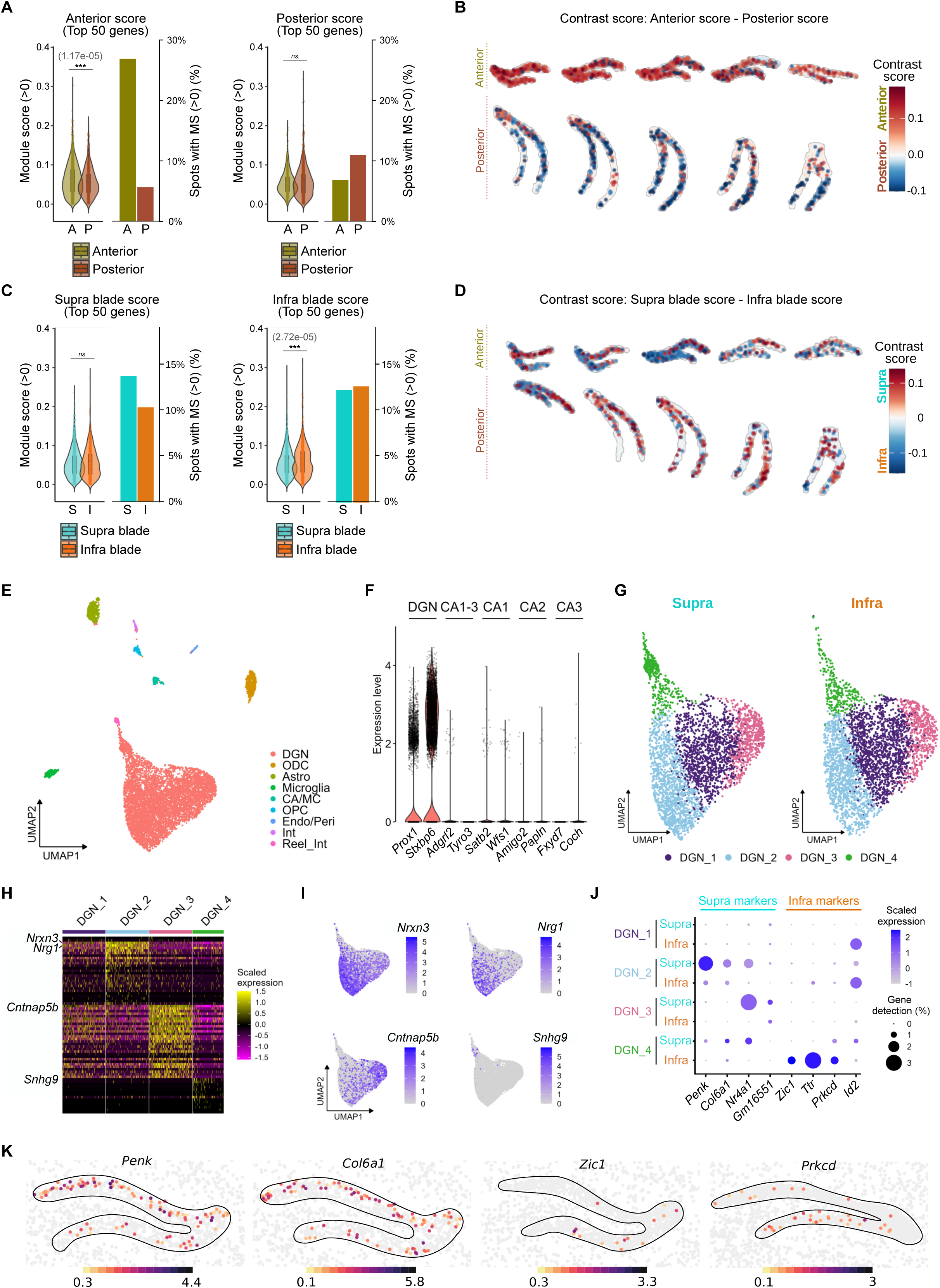
Spatial and single-nucleus transcriptomics reveal transverse molecular gradients in the DG. **(A)** Spatial module scores derived from bulk RNA-seq signatures. Violin plots show module scores across spots in anterior (A) and posterior (P) regions, and bar plots show the proportion of spots with positive module scores (>0) in each region. Scores were computed using the top 50 differentially expressed genes (FDR < 0.05) ranked by log_2_ fold change. Left panels correspond to anterior-enriched signatures and right panels to posterior-enriched signatures. **(B)** Spatial distribution of contrast scores (Score = Score_up − Score_down) for anterior vs posterior transcriptional signatures across representative Slide-seq pucks. Five anterior (top) and five posterior (bottom) pucks are shown. Positive scores (red) indicate anterior gene enrichment, whereas negative scores (blue) indicate posterior enrichment. **(C)** As in (A), violin plots show the distributions in supra (S) and infra (I) blades, and bar plots show the fraction of spots with positive scores (>0) in each blade. Scores were computed using suprapyramidal- (left) and infrapyramidal-enriched (right) bulk RNA-seq signatures. **(D)** As in (B), showing contrast scores for suprapyramidal vs infrapyramidal across five anterior (top) and five posterior (bottom) pucks. Positive scores (red) indicate suprapyramidal gene enrichment, whereas negative scores (blue) indicate infrapyramidal enrichment. **(E)** UMAP plots of DG nuclei. Nine main clusters were identified, including the most common cell types in this brain area: Astro: Astrocytes; ODC: Oligodendrocytes; CA/MC: Cornu Ammonis/Mossy Cells; OPC: Oligodendrocyte Progenitor Cells; Endo/Peri: Endocytes/Pericytes; Int: Interneurons; Reel_Int: Reelin-expressing interneurons. **(F)** Violin plot showing the expression of DGN markers but not CA markers in the clusters corresponding to DGN nuclei. **(G)** UMAP plots of the four DGN subclusters identified in the analysis. The clusters were present in the two blades. **(H)** Heatmap showing normalized expression of the markers of each subcluster of DGNs. Note the presence of a small number of markers of DGN_1 nuclei in comparison to the other subclusters. **(I)** Expression of four markers of subclusters DGN_2, DGN_3 and DGN_4. With the exception of *Snhg9* (which is exclusively expressed in DGN_4 nuclei), the other markers show a gradient expression pattern. **(J)** Dot plot of the expression of blade-enriched markers in DGN subclusters. **(K)** MERFISH images obtained from the Brain Knowledge Platform (https://knowledge.brain-map.org/abcatlas)^55^ illustrating the preferential expression of two suprapyramidal DGN markers (*Penk* and *Col6a1*) and two infrapyramidal DGN markers (*Zic1* and *Prkcd*) identified in bulk RNA-seq. Color bars indicate the minimum and maximum expression level as Log_2_(Counts Per Million + 1) for each image.

**Figure 3:**
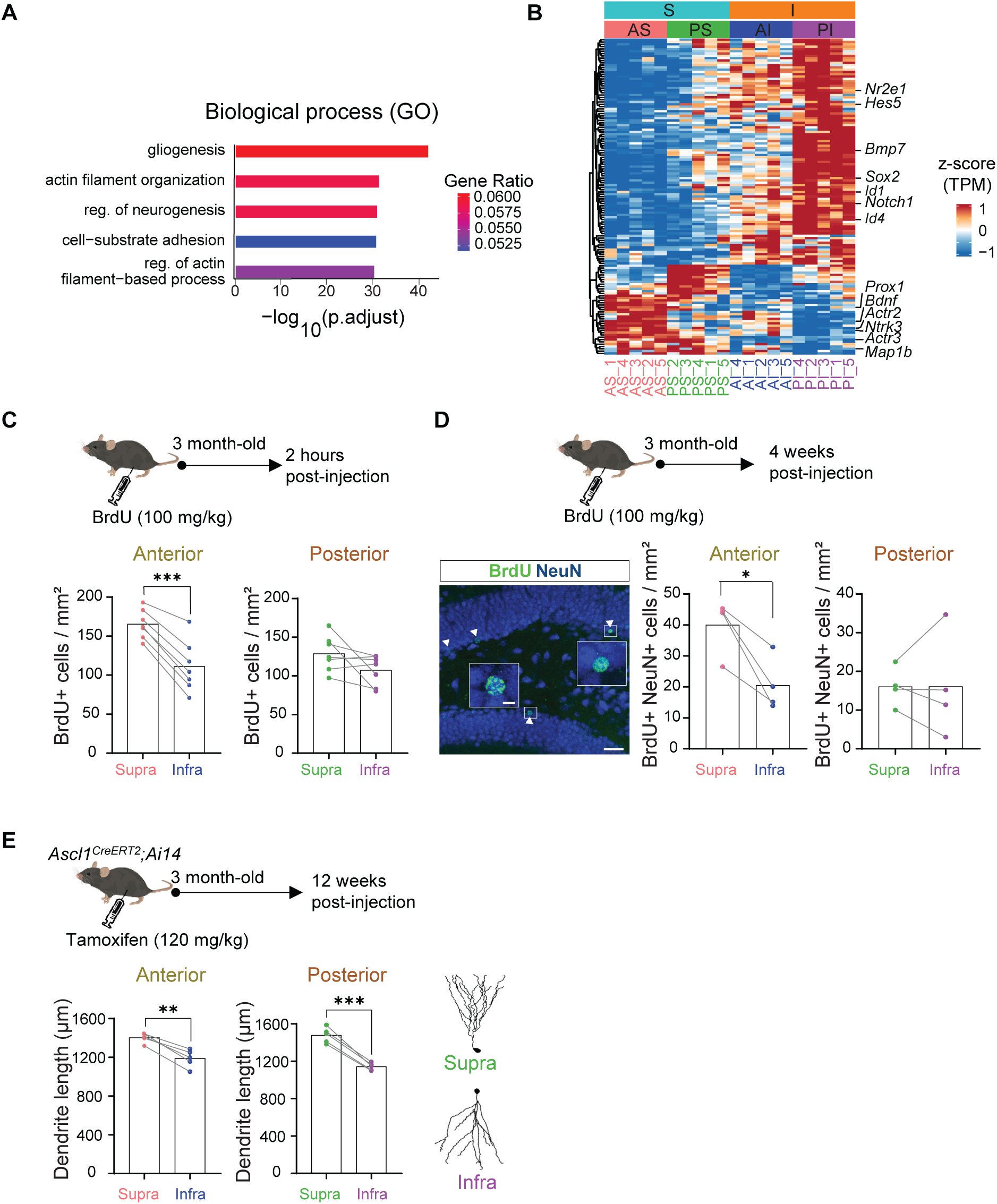
Adult neurogenesis along the transverse axis of the DG. **(A)** GO enrichment analysis of biological processes associated with SB vs IB DEGs. The top five terms (by FDR) are shown; bar color indicates gene ratio. **(B)** Heatmap of genes associated with the GO term “regulation of neurogenesis” from the SB vs IB comparison (row z-scored mean TPM values). **(C)** Quantification of BrdU⁺ cells/mm^2^ in the SB and IB of the anterior and posterior DG, 2h after BrdU injection in adult animals. Mean ± s.e.m., paired two-tailed Student’s t-test; \*\*\**p*<0.001 (n=7 mice). **(D)** Quantification of BrdU⁺ NeuN+ cells/mm^2^ in the SB and IB of the anterior and posterior DG, 4 weeks after BrdU injection in adult animals. Mean ± s.e.m., paired two-tailed Student’s t-test; \**p*-value<0.05 (n=4 mice). Scale bars: 50 µm and 5 µm in insets. **(E)** Quantification of total dendritic length of adult-born DGNs in the SB and IB of the anterior and posterior DG. Representative reconstructions of DGNs in the posterior DG. Mean ± s.e.m., paired two-tailed Student’s t-test; \*\**p*<0.01, \*\*\**p*<0.001 (n=5 mice).

## RESULTS

### Dentate gyrus blades are molecularly distinct compartments

To determine whether DG blades differ in molecular identity, we profiled the mRNA transcriptome of microdissected SB and IB of the adult mouse DG using bulk RNA-seq (Figure 1A). We analyzed blade transcriptomes separately in anterior and posterior DG, resulting in four subregions: anterior suprapyramidal (AS), anterior infrapyramidal (AI), posterior suprapyramidal (PS) and posterior infrapyramidal (PI) blades (Figure 1A; see methods). Replicates from each microdissected subregion showed strong correlation (*r* > 0.99; Figure S1A), indicating high reproducibility and internal consistency of the dataset. Analysis of established regional markers^49–53^ confirmed enrichment of DG markers (*Prox1, Igfbp5, Stxbp6)* and negligible expression of cornu ammonis markers CA1-3 *(Adgrl2*, *Tyro3*), CA1 *(Satb2, Wfs1,Pex5l*), CA2 (*Amigo2, Papln, Dcn, Cacng5*), CA3 (*Bok, Fxyd7*) (Figure S1B), demonstrating the specificity of our microdissection and dataset.

Hierarchical clustering of all samples revealed clear segregation according to DG subregion (Figure 1B). The primary separation distinguished anterior (A) and posterior (P) samples. Within each branch, samples further segregated into supra (S) and infra (I) subgroups (Figure 1B). Principal component analysis (PCA) similarly resolved four distinct and non-overlapping clusters corresponding to the four DG subregions (Figures 1C and S1C), supporting that these regions are molecularly distinct.

Differential gene expression analysis revealed pronounced transcriptional differences among DG subregions. As expected, many genes exhibited anteroposterior expression differences across the DG (Figures S1D-E), and similar regional differences were observed within each blade (Figures S1F-G; Table S1). Importantly, we also identified robust blade-specific gene expression differences (Figure S1H), with distinct transcriptional patterns in anterior and posterior DG (Figures 1D-G, Table S1). In the anterior DG, 900 genes were downregulated and 397 upregulated in the SB versus IB (Figure 1D), whereas in the posterior DG, 805 genes were downregulated and 476 upregulated (FDR ≤ 0.05) (Figure 1E). Notably, several genes showed strong downregulation (log_2_FC < −3, FDR ≤ 0.01) in AS versus AI and PS versus PI comparisons, identifying potential regional markers for AI or PI, such as *Ttr* and *Tnnt1* respectively (Figure 1G). Although no genes reached log_2_FC > 3, multiple genes showed moderate but significant upregulation (log_2_FC > 1, FDR < 0.01) and were selectively enriched in AS or PS, such as *Col6a1* and *Cartpt* (Figure 1G). Together, these results indicate that both blade identity and anterior–posterior position contribute to transcriptional heterogeneity within the DG.

To validate blade-specific gene expression patterns, we examined selected differentially expressed genes (DEGs) using qPCR, *in situ* hybridization data from the Allen Brain Atlas^54^, and publicly available MERFISH datasets^55^ (Figure S2A-D). SB-enriched genes such as *Penk*, *Col6a1*, and *Cartpt* showed expression patterns consistent with RNA-seq data across datasets, with *Penk* enriched in both anterior and posterior SB, *Col6a1* selectively enriched in anterior SB, and *Cartpt* enriched in posterior SB (Figure S2A-D). Similarly, the expression patterns of the IB-enriched genes, *Zic1, Ttr, Tnnt1*, were confirmed (Figure S2A,B,D). Together, these cross-platform validations support the robustness of our dataset and support the reliability of blade-specific gene expression patterns. We made this dataset publicly available as the DG-seq resource (https://dgseq.neurocentre-magendie.fr/).

To identify gene expression programs associated with DG subregions, we grouped genes into co-expression modules using network analysis^35^. This analysis identified 40 gene co-expression modules, of which 11 showed significant differences across DG subregions (Figure S3, Table S2). Correlation of module eigengenes (MEs) with DG subregions revealed strong association between specific modules and individual blades or regions (|R| ≥ 0.5, *p* -value ≤ 0.05) (Figure 1H). Examination of module expression patterns across DG subregions revealed distinct transverse-axis transcriptional programs. For example, module _22 was enriched in SB across both anterior and posterior regions, whereas module_9 was enriched specifically in the anterior DG (Figure 1I). Similarly, module_1 was enriched in IB across regions, while module_*5* was enriched specifically in the anterior IB (Figure 1J).

To gain insights into the functional implication of these modules, we performed Gene Ontology (GO) enrichment analysis, which revealed recurrent enrichment of neurogenesis related biological processes across multiple modules (Figure 1K, Table S3), suggesting functional differences in neurogenic regulation across DG subregions. Together, these results demonstrate that the SB and IB DG blades constitute molecularly distinct compartments with anterior-posterior gene expression programs, supporting the idea that DG subregions are molecularly and functionally specialized.

### Spatial transcriptomics reveals robust spatial organization of DG transcriptional programs

To determine whether the transcriptional differences identified by anatomically resolved bulk RNA-seq were spatially organized within the DG, we integrated these data with a Slide-seq dataset spanning the entire adult mouse hippocampus (Figure S4A)^41^. Given the limited transcript detection sensitivity of Slide-seq, spatial spots across all pucks were scored using gene signatures derived from the bulk RNA-seq analysis. Spots located in the anterior DG exhibited significantly higher module scores for the anterior gene signature than those in the posterior DG (Figure 2A). In addition, anterior signature-positive spots (module score > 0) were substantially more frequent in the anterior DG, indicating both stronger and more widespread representation of the anterior transcriptional program in this region (Figure 2A). In contrast, posterior gene signature scores showed only a trend towards higher values in posterior DG spots (Figure 2A). However, posterior signature-positive spots were more abundant in the posterior DG than in the anterior DG, suggesting that enrichment of the posterior transcriptional program is primarily driven by a larger proportion of signature-positive spots rather than by stronger expression within individual spots (Figure 2A). To further assess the spatial segregation of these transcriptional programs, we calculated a contrast score by subtracting the posterior signature score from the anterior signature score for each spot. This analysis revealed a striking spatial segregation of these signatures, with anterior-enriched transcriptional programs localizing predominantly to the anterior DG and posterior-enriched programs to the posterior DG (Figure 2B). Notably, this anterior–posterior organization was independently recapitulated in both the suprapyramidal and infrapyramidal blades (Figure S4B–D). Altogether, these findings demonstrate that the DG is organized into spatially distinct transcriptional domains along its longitudinal axis.

We next asked whether transcriptional differences between the SB and IB identified by bulk RNA-seq were similarly preserved in the spatial transcriptomic dataset. Module scores for the suprapyramidal gene signature were comparable between blades (Figure 2C). However, suprapyramidal signature-positive spots (module score > 0) were substantially more frequent in the SB, indicating a broader spatial representation of the suprapyramidal transcriptional program in this region. Conversely, infrapyramidal gene signature scores were significantly higher in IB spots than in SB spots. In contrast to the suprapyramidal signature, the proportion of infrapyramidal signature-positive spots was similar between blades, suggesting that enrichment of the infrapyramidal transcriptional program is primarily driven by stronger expression within individual spots rather than by an increased spatial prevalence of positive spots (Figure 2C). To further assess the spatial segregation of blade-specific transcriptional programs, we calculated a contrast score by subtracting the infrapyramidal signature score from the suprapyramidal signature score for each spot. This analysis revealed a clear segregation of transcriptional identities across the transverse axis of the DG, with suprapyramidal- and infrapyramidal-enriched programs localizing to their respective blades (Figure 2D). Importantly, this blade-specific organization was maintained throughout the anterior–posterior extent of the DG and was independently recapitulated when anterior and posterior DG regions were analysed separately (Figures S4E–G). Altogether, these findings demonstrate that the molecular specialization of the SB and IB is preserved along the longitudinal axis of the DG.

### Single-nucleus RNA sequencing reveals cell-type contributions to blade-specific gene expression

To determine the cellular origin of blade-specific gene expression differences, we performed single-nucleus RNA sequencing (snRNA-seq) of nuclei isolated from the anterior DG (Figure S5A–C). After quality control filtering, 8,414 nuclei were recovered and clustered into nine major cell populations corresponding to the principal DG cell types based on established marker genes (Figure 2E, Table S4). Dentate granule neurons (DGNs), expressing *Prox1* and *Stxbp6* but not CA pyramidal cell markers (Figure 2F), formed four subclusters (Figure 2G). Non-linear dimensionality reduction showed that the four DGNs clusters were tightly packed, indicating the presence of gene expression gradients, rather than sharp differences in gene expression between clusters, with all DGNs clusters preserved across SB and IB (Figure 2G). The top markers of each cluster were identified, which revealed the presence of three well-defined clusters (DGN_2, DGN_3 and DGN_4) and one less differentiated cluster (DGN_1) (Figure 2H, Table S5). As expected, the main markers of the more differentiated clusters exhibited gradient expression patterns (Figure 2I).

We then examined the expression of blade-enriched genes identified in the bulk RNA-seq dataset across these cell clusters (Table S6). SB-enriched genes such as *Penk, Col6a* and *Nr4a1* were primarily expressed in DGNs, with a higher number of expressing cells in the SB compared to the IB (Figures 2J and S5D–E), consistent with the bulk RNA-seq results. In contrast, IB-enriched genes such as *Id2*, *Zic1, Ttr* and *Prkcd* were expressed across multiple cell types but were consistently detected in a higher number of IB cells or were restricted to IB cells (Figures 2J and S5D–E). Within DGNs, blade-enriched genes showed variable expression across the four DGN clusters (Figure 2J). Blade-specific marker expression was further validated using publicly available single-cell transcriptomic data from the Allen Brain Cell Atlas^55^, focusing on glutamatergic DGNs (Figure 2K).

Together, these results indicate that blade-enriched genes are not confined to blade-specific cell clusters but instead are differentially expressed across cell populations shared between blades, supporting the molecular specialization of DG blades.

### Molecular blade identity reveals difference in adult neurogenesis along the transverse axis

To assess the functional implications of blade-specific gene expression differences, we performed Gene Ontology enrichment analysis of DEGs between the SB and IB identified in our bulk RNA-seq data (Figure S1H). This analysis revealed significant enrichment of biological processes related to gliogenesis, actin filament organization, and regulation of neurogenesis (adjusted *p*-value < 0.05; Figure 3A). Consistent with our previous module-based GO analysis (Figure 1K), genes associated with regulation of neurogenesis showed distinct expression patterns between the two blades in both anterior and posterior regions (Figure 3B, Table S7). These findings suggest blade-specific neurogenic programs with additional regional variation along the anterior–posterior axis of the adult DG. Notably, the identified genes were associated with multiple stages of neurogenesis, including proliferation (e.g. *Sox2, Id1*), neuronal differentiation and fate commitment (e.g. *Ntrk3*, *Prox1*), and neuronal maturation such as dendritogenesis (e.g. *Actr3*, *Bdnf)*.

We next experimentally assessed whether these adult neurogenesis-related processes differ between the two DG blades. To examine proliferation, 3-month-old mice received a single intraperitoneal injection of the thymidine analog BrdU (100 mg/kg) and were sacrificed 2h later. Quantification of BrdU⁺ cells revealed higher proliferation in SB compared to the IB in the anterior DG (165.5 ± 7.0 vs 111.1 ± 12.4 cells/mm^2^, t_6_=9.54, *p*<0.0001), whereas no significant differences were observed in the posterior DG (128.8 ± 8.7 vs 107.4 ± 7.3 cells/mm^2^, t_6_=1.89, *p*=0.11) (Figure 3C). To assess neuronal differentiation, animals were sacrificed four weeks after BrdU injection to quantify BrdU+ cells co-expressing the mature neuronal marker NeuN. Neuron generation was higher in the SB compared to IB in the anterior DG (39.1 ± 4.3 vs 20.5 ± 4.4 cells/mm^2^, t_3_=4.30, *p*=0.02), whereas no differences were detected between blades in the posterior DG (16.0 ± 2.6 vs 16.1 ± 6.7 cells/mm^2^, t_3_=0.006, *p*=1.00) (Figure 3D). Finally, to evaluate dendritic maturation of adult-born neurons, we labelled newborn neurons by injecting tamoxifen to 3-month-old *Ascl1^CreERT2^;Ai14 mice* and analyzed dendritic morphology 12 weeks later. Total dendritic length was significantly greater in the SB compared to IB in both anterior (1402.2 ± 22.5 vs 1188.7 ± 40.7 µm, t_4_=7.58, *p*=0.002) and posterior DG (1478.4 ± 37.1 vs 1143.0 ± 18.0 µm, t_4_=12.43, *p*<0.001) (Figure 3E).

Altogether, these results demonstrate that adult neurogenesis is differentially regulated between DG blades, with higher neurogenic activity in the SB and region-specific differences along the anterior–posterior axis.

### The IB contains a larger pool of quiescent neural stem cells

Given the differences in adult neurogenesis between blades, we next asked whether neural stem cell (NSC) populations and quiescence states differ between the SB and IB. To address this, we performed cell-type marker enrichment analysis using over-representation analysis of DEGs between the two blades. This analysis revealed a significant overrepresentation (adjusted *p*- value < 0.05) of markers associated with NSCs, including quiescent NSCs (qNSC) (Figure 4A). Consistently, analysis of previously reported NSC quiescence markers^56–59^ showed that most were expressed at higher levels in the IB compared to the SB across DG subregions (Figure 4B). MERFISH data^55^ further confirmed higher expression of *Hopx*, *GFAP*, and *Aqp4*, genes associated with NSC quiescence, in the IB relative to the SB in both anterior and posterior DG (Figure 4C).

**Figure 4:**
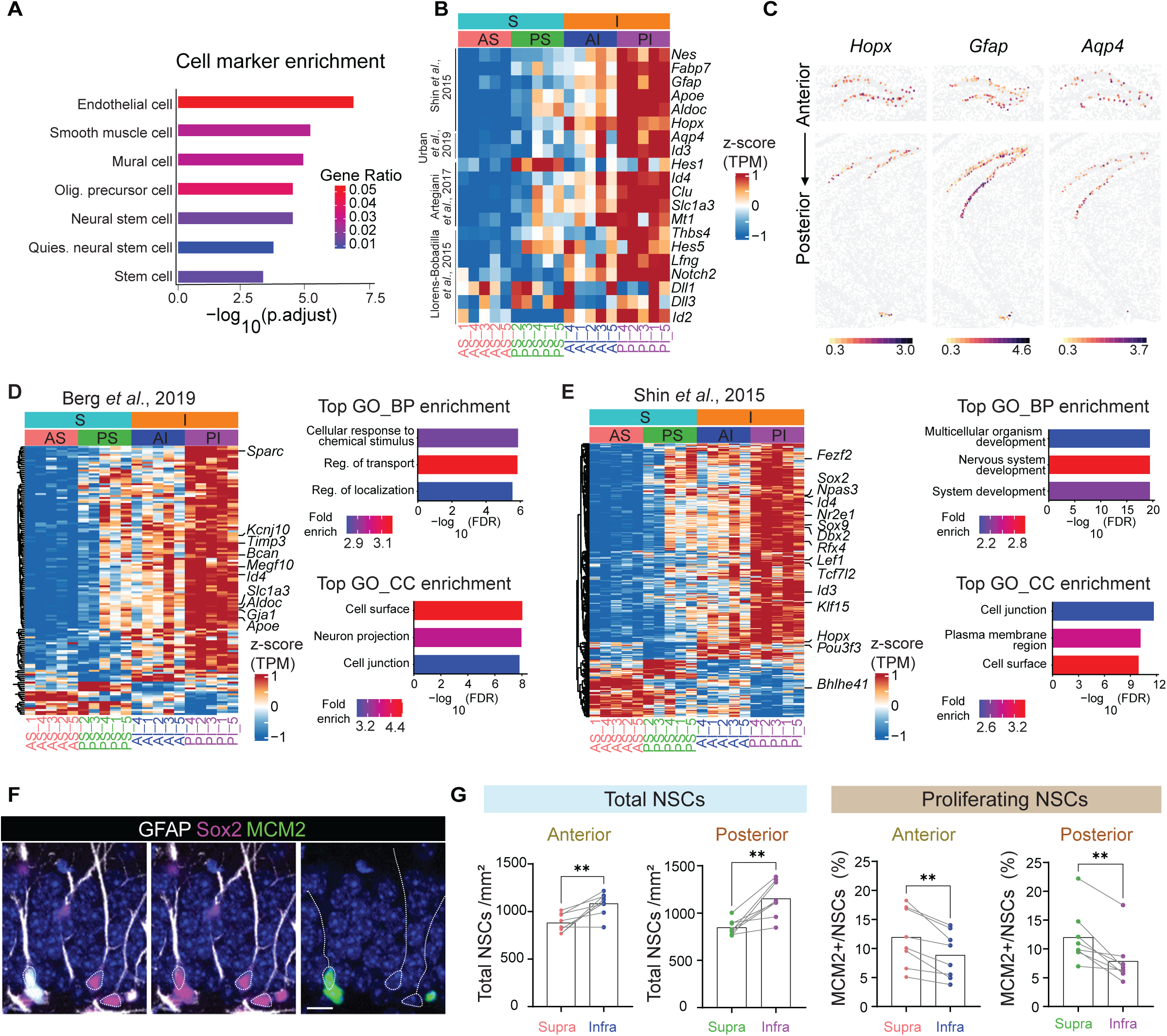
Adult NSC quiescence along the transverse axis of the DG. **(A)** Over-representation analysis of cell-type markers among SB vs IB DEGs (CellMarker 2.0, Mus musculus brain). Top terms ranked by FDR; bar color indicates gene ratio. **(B)** Heatmap of reported quiescent NSC marker genes detected in our dataset (row z-scored mean TPM values). **(C)** Expression of selected NSC quiescence genes in the DG from MERFISH data^62^. Parameters for selection: Zhuang-ABCA-1, Cell properties, Allen Mouse Common Coordinate “DG-sg”. Color bars indicate the minimum and maximum expression level as Log_2_(Counts Per Million + 1) for each image. **(D–E)** Heatmaps of DEGs overlapping with published quiescent NSC gene signatures^58,60^, shown as row z-scored mean TPM values. **(G)** Representative images of NSCs (GFAP+ Sox2+ radial glia-like cells) and proliferating NSCs (GFAP+, Sox2+ and MCM2+ radial glia-like cells). Scale bar: 10 µm. **(H)** Quantification of total NSC/mm^2^ and proportion of proliferating NSC in SB and IB of anterior and posterior DG. Mean ± s.e.m, paired two-tailed Student’s t-test; \*\**p*<0.01, n=8 mice.

To further characterize blade-specific molecular differences related to stem cell quiescence, we compared our DEGs with published qNSC gene signatures^58,60^. Of the 283 qNSC-associated genes reported by Berg et al.^60^, 137 overlapped with our DEGs and were mainly enriched in the IB (Figure 4D). Similarly, 314 of 1000 qNSC-associated genes reported by Shin et al.^58^ overlapped with our DEGs (Figure 4E). Among these, transcription factors previously linked to qNSC identity, including *Sox2*, *Sox9*, *Id3*, and *Nr2e1*, also showed higher expression in the IB (Figure 4E). Functional enrichment analysis of the overlapping genes identified pathways related to cellular response to chemical stimulus, molecular transport, and cell-surface signaling genes, consistent with previously described qNSC molecular signatures^58,60^.

These findings suggesting blade-specific differences in NSC quiescence were further examined using immunohistochemistry. We first quantified total NSCs, identified as GFAP+ cells with radial-glia morphology co-expressing Sox2 in the SGZ (Figure 4F). A higher density of NSCs was observed in the IB compared to the SB in both anterior (1081.8 ± 42.9 vs 879.8 ± 34.5 cells/mm^2^, t_7_=4.03, *p*=0.005) and posterior DG (1151.6 ± 67.9 vs 846.0 ± 28.4 cells/mm^2^, t_7_=4.59, *p*=0.003) (Figure 4G). Moreover, MCM2 staining revealed a significantly lower proportion of proliferating NSCs in the IB across the entire DG (anterior: 11.9 ± 1.8% in SB vs 8.9 ± 1.4% in IB, t_7_=4.51, *p*=0.003; posterior: 12.0 ± 1.7% in SB vs 7.9 ± 1.5% in IB, t_7_=4.68, *p*=0.002), indicating a larger population of quiescent NSCs in this blade (Figure 4G). Together, these findings indicate that the IB harbors a larger reservoir of quiescent NSCs, whereas the SB exhibits higher neurogenic activity.

## DISCUSSION

In this study, we demonstrate that the SB and IB of the adult DG constitute molecularly and functionally distinct hippocampal compartments rather than simple anatomical subdivisions. Through the integration of bulk RNA-seq from four anatomically defined DG subregions— anterior SB, anterior IB, posterior SB, and posterior IB—with spatial transcriptomics, snRNA-seq, and functional analyses, we uncover a previously underappreciated axis of heterogeneity within the DG. Importantly, blade-specific signatures are not uniform across the DG but vary along the anterior–posterior axis, revealing an interaction between transverse and longitudinal patterning in shaping DG identity. Extending previous work on dorsoventral organization^49^, these findings establish the DG as a fundamentally multi-axial structure and underscore the importance of considering multiple anatomical dimensions when studying DG organization and function in health and disease.

A striking finding of this study is that SB and IB identity is encoded by transcriptional programs that are consistently detected across bulk RNA-seq, spatial transcriptomics, snRNA-seq, and independent datasets, indicating that blade identity is a fundamental and robust organizational feature of the DG. The spatial recovery of these signatures further demonstrates that blade-specific molecular programs are organized into anatomically defined domains across the adult DG. These programs include IB-enriched genes such as *Zic1*, *Ttr*, and *Tnnt1*, and SB- enriched genes such as *Penk*, *Col6a1*, and *Cartpt*, whose expression patterns are either broadly distributed across blades or restricted to specific anterior or posterior subregions. Although most of these genes are reported here for the first time as blade-enriched, *Penk* was primarily expressed in the DGN_2 cluster and showed preferential expression in the SB blade, in agreement with previous studies^22,61^. Some of these genes are expressed in relatively sparse cell populations; nevertheless, their reproducible blade-specific enrichment suggests that rare cellular states may contribute disproportionately to regional identity. Accordingly, blade- enriched transcriptional programs may have outsized effects on circuit function and information processing. Future studies will be required to determine how these molecular signatures contribute to blade-specific circuit dynamics and behavioral function.

In addition to regional molecular heterogeneity, we reveal cellular diversity within the DGN population. Although DGNs have traditionally been considered relatively homogeneous, recent snRNA-seq studies in mice and humans have identified multiple transcriptionally distinct DGN subclusters^22,62^. Consistently, our analysis uncovers four DGN subclusters characterized by graded transcriptional differences. Whether these subclusters reflect DGNs generated during distinct developmental periods^48,63^, different stages of neuronal maturation, various functional states, or a combination of these factors remains unclear. Notably, each subcluster exhibits distinct gene expression between the SB and IB blades, indicating that blade identity intersects with intrinsic DGN heterogeneity. Such diversity likely contributes to functional specialization within the DG and may provide a cellular substrate for the computational flexibility of hippocampal circuits^64^, enabling a broader repertoire of neural computations and behavioral outputs. Understanding how these molecularly defined DGN subpopulations within each blade contribute to circuit function and behavior remains an important challenge for future studies.

Functional analyses indicate that blade-specific molecular programs are associated with distinct regulation of adult neurogenesis. Although previous studies have reported blade-specific patterns of adult neurogenesis under basal conditions, their findings have been inconsistent^65–68^. By resolving these discrepancies along the anterior–posterior axis in adult mice, we demonstrate that blade-dependent neurogenesis is not uniform throughout the DG but is instead more pronounced in the anterior DG. Specifically, we observed increased neurogenic activity, including progenitor proliferation and newborn neuron generation, in the anterior SB. In contrast, the IB harbors a larger pool of quiescent NSCs, providing the first molecular evidence that stem cell states are spatially compartmentalized across DG blades. Given that adult qNSCs actively integrate diverse niche-derived signals^58^, these findings raise the possibility that IB NSCs are tuned to distinct extrinsic cues compared with their SB counterparts, reflecting blade-specific microenvironmental and circuit-level regulation of stem cell behavior. Collectively, our data support a model in which the SB is biased toward active neurogenesis and circuit remodeling, whereas the IB functions as a reservoir of quiescent NSCs that can be mobilized in response to specific physiological or pathological demands.

Together, these findings establish the DG as a multidimensional structure in which transverse (SB–IB) and longitudinal (anterior–posterior) axes jointly shape molecular identity, cellular composition, and neurogenic dynamics, rather than serving as a uniform gateway to the hippocampus. This framework uncovers spatially organized stem cell states and blade-specific molecular programs that differentially regulate adult neurogenesis, highlighting functional specialization across DG compartments. By providing a comprehensive molecular map of DG organization, our study lays the foundation for understanding how spatial heterogeneity within the DG contributes to hippocampal circuit function in health and disease.

## RESOURCE AVAILABILITY

### Lead contact

Further information should be directed to and will be fulfilled by the lead contact, Emilie Pacary

## Materials availability

This study did not generate any new unique reagents.

## Data and code availability

Raw fastq files and counts matrices are available on Gene Expression Omnibus (GEO) database: **<u>GSE313634</u>** (bulk RNA-seq), **<u>GSE311291</u>**(snRNA-seq) are grouped under SuperSeries **<u>GSE313937</u>**. A web application provides visualization of bulk RNA-seq data (DG-seq): https://dgseq.neurocentre-magendie.fr/

## ACKNOWLEDGMENTS

This work was supported by Inserm, ANR (ANR-19-CE16-0014-01 and ANR-25-CE16-6626- 01 to E.P.), FRM (Fondation pour la Recherche Medicale, EQU202203014657 to D.N.A.) and Neuroglia (grant to E.P.). This work was supported by MICIU/AEI /10.13039/501100011033 and ERFD, EU (grants PID2021-129053OB-I00 and PID2024-163108OB-I00 to J.P.L-A; grant PID2021-124829NB-I00 to L.M.d.l.P.). J.P.L-A. research is supported by grant CIPROM/2023/15 funded by the Generalitat Valenciana, Conselleria d’Educació, Universitats i Ocupació. J.P.L-A. acknowledges financial support from Centro de Excelencia Severo Ochoa, grant CEX2021-001165-S, funded by MCIN/AEI/10.13039/501100011033. P.T.M. was supported by Fundación Tatiana and a FENS/IBRO-PERC Exchange Fellowship for a three-month stay in E.P.’s lab. This work benefited from the support of the Microdissection Laser Capture facility funded by Inserm, LabEX BRAIN ANR-10-LABX-43 and FRM DGE20061007758, the Transcriptomic facility funded by Inserm and LabEX BRAIN ANR-10- LABX-43 and the Animal Housing and Genotyping facilities funded by Inserm and LabEX BRAIN ANR-10-LABX-43. RNA sequencing was performed at the PGTB (https://doi.org/10.15454/1.5572396583599417E12) with the help of Zoé Compagnie. Acquisitions with the confocal Leica SP5 or SP8 were done in the Bordeaux Imaging Center (BIC), a service unit of the CNRS-Inserm and Bordeaux University, member of the national infrastructure France BioImaging supported by the French National Research Agency (ANR- 10-INBS-04). The authors thank Antonio Caler for technical help in the FACS/Omics facility of the Instituto de Neurociencias (UMH-CSIC). We are grateful to Jaime Pignatelli and Aura Torres Medina at the Centro de Neurociencias Cajal (CNC, CSIC) for exploratory spatial transcriptomics analyses and for valuable discussions.

## AUTHOR CONTRIBUTIONS

U.A.B., E.P., and J.P.L-A. conceptualized the study and designed the experiments.

U.A.B. analyzed bulk RNA-seq data, developed the DG-seq website, prepared brain samples, and performed dendrite analyses.

A.M.G. analyzed spatial transcriptomics data.

C.M.N. dissected DG blades, isolated DG cells and prepared snRNA-seq libraries.

S.V. and A.M.G. analyzed snRNA-seq data.

P.T.M. performed NSC immunohistochemistry and analysis under the supervision of A.V.M.

M.V. performed BrdU immunostaining for proliferation and conducted the analysis.

F.M. prepared the libraries.

H.D. and M.M. microdissected the blades for bulk RNA-seq.

V.G. and T.L.S. extracted RNA and performed RT-PCR.

U.A.B. and A.M.G. performed data curation and completed the GEO submission.

A.B. contributed to website hosting.

U.A.B., S.V., A.M.G., D.N.A., L.M.d.l.P., J.P.L-A., and E.P. discussed the results and wrote the manuscript.

E.P., D.N.A., L.M.d.l.P. and J.P.L-A. provided funding.

EP and J.P.L-A. supervised the entire project.

