## Supplementary information for "Blade-dependent molecular identity and neurogenic potential in the adult dentate gyrus"

### SUPPLEMENTARY FIGURE LEGENDS

**Figure S1: Quality control and reproducibility of the DG bulk RNA-seq.**

**(A)** Pearson correlation matrix across all samples, with colors ranging from blue to red indicating increasing levels of correlation among samples.

**(B)** Heatmap visualizing the expression profiles of reported marker genes across samples and DG subregions, based on column-scaled TPM counts.

**(C)** PCA biplot showing the distribution of samples and gene loadings for a selected set of genes. Small points represent individual samples, while larger points indicate group means. The length and direction of the gene loading arrows reflect the strength and direction of each gene's association with the principal components.

**(D)** Volcano plots illustrating gene expression changes between the posterior DG and anterior DG. Samples from AS and AI were digitally pooled as anterior DG, and PS and PI as posterior DG. RNAs with a false discovery rate (FDR) $\leq 0.05$ were considered significant and are shown in red (upregulated genes; 1624 genes) and blue (downregulated; 1848 genes) in the anterior DG relative to the posterior DG.

**(E)** Heatmap based on row-scaled TPM values showing the comparative expression profiles of top genes exhibiting anterior–posterior expression polarity in the bulk RNA-seq, compared with the reported dorsal–ventral expression profiles of the same genes in Cembrowski et al., 2016 [49]. Note that the posterior sample in the bulk RNA-seq includes most of the ventral DG.

**(F-G)** Volcano plots illustrating differential gene expression between posterior and anterior SB (F) and between posterior and anterior IB (G).

**(H)** Volcano plots illustrating gene expression differences between the SB and IB across the DG (999 upregulated and 1482 downregulated genes, FDR $\leq 0.05$). Samples from AS and PS were pooled as SB, and those from AI and PI as IB.

**Figure S2: Orthogonal cross-platform validation of selected genes.**

**(A)** Expression (TPM counts) profiles of selected genes across DG subregions.

**(B)** Comparison of expression profiles of selected genes measured by RNA-seq and qPCR. Pearson correlations were calculated for each gene in a subregion-specific manner between $\log_2$-transformed TPM values and $\log_2$ fold-change between blades. Statistical significance was assessed using a permutation test.

**(C)** Expression profiles of selected genes in the DG, derived from the Allen Brain Atlas (ABA). Scale bars represent 500 μm.

**(D)** MERFISH images of selected genes obtained from the Brain Knowledge Platform (https://knowledge.brain-map.org/abcatlas)[55]. Parameters used for selection: Zhuang-ABCA-1, Cell properties, Allen Mouse Common Coordinate "DG-sg". *Ttr* and *Tnnt1* were not detected in ABA or MERFISH, likely due to low expression levels. Color bars indicate the minimum and maximum expression level as $Log_2$(Counts Per Million + 1) for each image.

**Figure S3: Gene module expression along the DG transverse axis.**
Box plots showing aggregated z-scores of expression across samples and DG subregions for all gene modules identified in the multiWGCNA analysis.

**Figure S4: Extended spatial analysis of transcriptomic gradients across all Slide-seq pucks.**
**(A)** Anatomical annotation of all Slide-seq pucks. Each spot is classified by anteroposterior position (A, P), blade identity (S, I), and their combined compartments (AS, AI, PS, PI).
**(B)** Spatial distribution of contrast scores (Contrast score = Score_up − Score_down) for anteroposterior transcriptional signatures derived from bulk RNA-seq across all pucks. Positive (red) and negative (blue) scores indicate anterior- or posterior-associated gene expression, respectively. Three contrasts are shown: from global (top), from suprapyramidal (middle), and from infrapyramidal (bottom).
**(C-D)** Spatial module scores for anteroposterior transcriptional signatures within each blade. As in Fig. 2A, violin plots show the distribution across DG spots, and bar plots show the proportion of spots with positive scores (>0). Gene sets were derived from bulk RNA-seq comparisons of anterior vs posterior within the SB (C) or IB (D), using the top 50 differentially expressed genes (FDR < 0.05) ranked by decreasing $log_2$ fold change. Scores are compared between anterior (A) and posterior (P) regions using unpaired two-sided Wilcoxon rank-sum tests.
**(E)** As in (B), spatial distribution of suprapyramidal vs infrapyramidal contrast scores across all pucks. Positive (red) and negative (blue) scores indicate supra- or infrapyramidal enrichment. Three contrasts are shown: from global (top), from anterior (middle), and from posterior (bottom).
**(F–G)** Spatial module scores for blade-specific signatures within anterior (F) or posterior (G) compartments. As in Fig. 2C, violin and bar plots show score distributions and proportions of positive spots (>0), respectively. Gene sets were derived from the top 50 differentially expressed genes (FDR < 0.05) in suprapyramidal vs infrapyramidal comparisons within anterior

(F) or posterior (G) compartments, using the top 50 differentially expressed genes (FDR < 0.05) ranked by decreasing $log_2$ fold change. Scores are compared between suprapyramidal (S) and infrapyramidal (I) blades using unpaired two-sided Wilcoxon rank-sum tests.

**Figure S5**: **Dissection and gating strategy for DG nuclei and expression of blade-enriched genes across non-DGN clusters.**

**(A)** Representative stereomicroscope images of endogenous tdTomato fluorescence in coronal hippocampal sections from an adult *Calb1-2A-dgCre*$^{+/-}$*;Ai9*$^{+/-}$ mouse before and after microdissection of either the suprapyramidal (bottom left) or infrapyramidal (bottom right) DG blade. From each hippocampus, only one blade (suprapyramidal or infrapyramidal) was dissected to ensure blade-specific sampling.

**(B-C)** Representative fluorescence-activated nuclei sorting (FANS) strategy for nuclei isolated from the suprapyramidal (B) and infrapyramidal (C) DG blades. Left, FSC-A versus SSC-A plot used to identify intact nuclei and exclude debris. Center, FSC-A versus FSC-H gating used to remove doublets. Right, DAPI fluorescence intensity plot used to select DAPI-positive nuclei for downstream analyses.

**(D)** Heatmap showing the normalized expression of canonical marker genes across all cell populations identified in the snRNA-seq dataset: Astro: Astrocytes; ODC: Oligodendrocytes; CA/MC: Cornu Ammonis/Mossy Cells; OPC: Oligodendrocyte Progenitor Cells; Endo/Peri: Endocytes/Pericytes; Int: Interneurons; Reel_Int: Reelin-expressing interneurons.

**(E)** Dot plot showing the expression of selected suprapyramidal- and infrapyramidal-enriched genes across non-DGN clusters. *Zic1* and *Ttr* are preferentially expressed in infrapyramidal endothelial-pericyte cells, whereas *Nr4a1*, *Id2* and *Prkcd* show similar expression patterns across corresponding suprapyramidal and infrapyramidal cell populations.

### SUPPLEMENTARY TABLE LEGENDS

**Table S1:** Differentially expressed genes in the bulk RNA-seq dataset, between DG subregions.

**Table S2:** Eigengenes for identified modules in DME (differential module analysis) using multiWGCNA.

**Table S3:** GO biological processes analysis results for the modules in Figure 1H.

**Table S4:** Differentially expressed marker genes for the major hippocampal cell populations identified in the snRNA-seq dataset.

**Table S5:** Differential gene expression analysis identifying enriched genes for dentate granule neuron (DGN) subpopulations in the snRNA-seq dataset.

**Table S6:** Differentially expressed genes between DG subregions in the snRNA-seq dataset. DEGs identified by comparing either all suprapyramidal vs all infrapyramidal DGNs combined or by comparing within single subpopulations (DGN_1-DGN_4) are pooled. The column "source" identifies the group tested.

**Table S7:** Differentially expressed genes in suprapyramidal (S) vs infrapyramidal blade (I) in the bulk RNA-seq dataset, related to the GO-term 'regulation of neurogenesis' in Figure 3B.

Figure S1

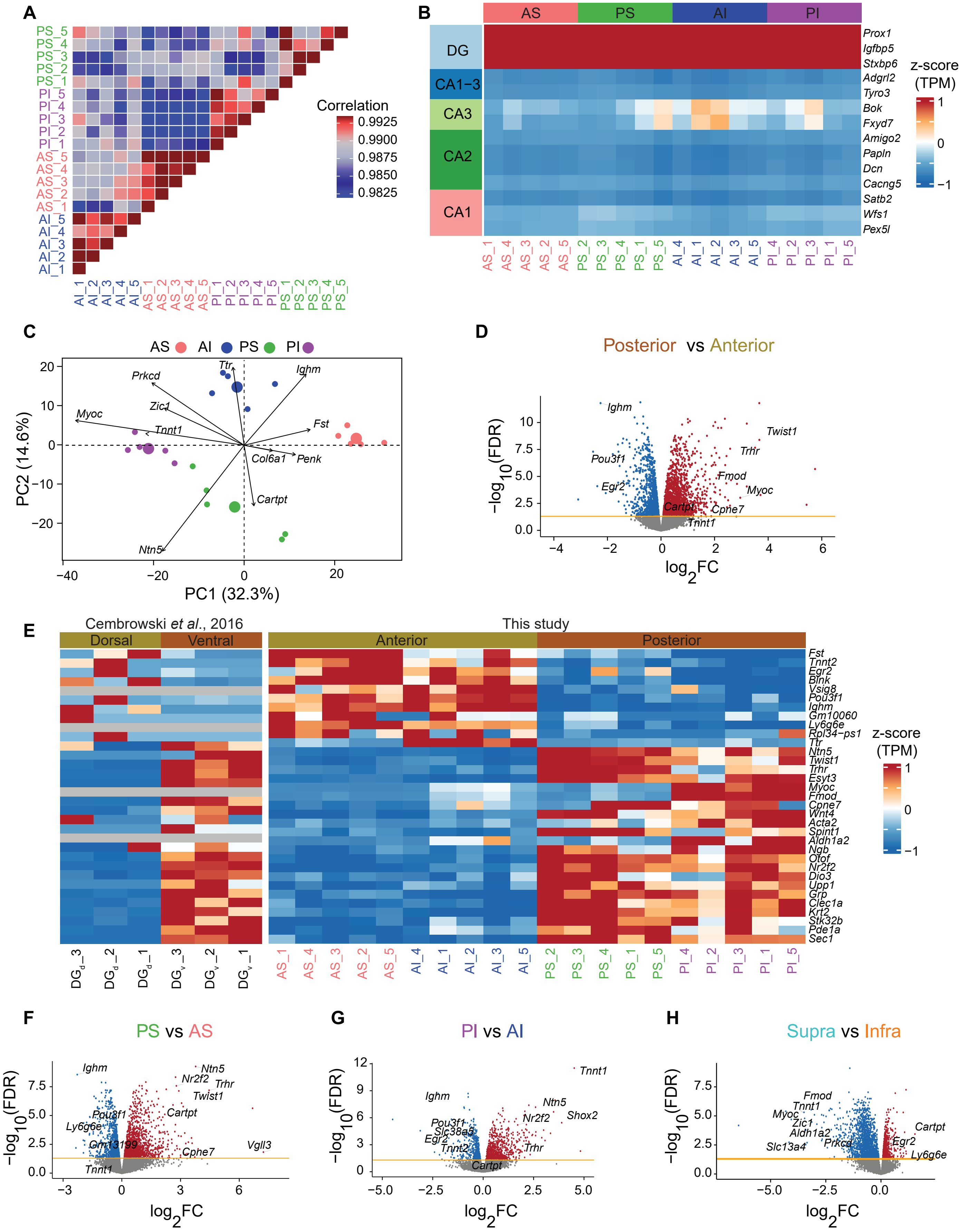
A
PS_5
PS_4
PS_3
PS_2
PS_1
PI_5
PI_4
PI_3
PI_2
PI_1
AS_5
AS_4
AS_3
AS_2
AS_1
AI_5
AI_4
AI_3
AI_2
AI_1
Correlation
0.9925
0.9900
0.9875
0.9850
0.9825
B
AS
PS
AI
PI
DG
CA1−3
CA3
CA2
CA1
Prox1
Igfbp5
Stxbp6
Adgrl2
Tyro3
Bok
Fxyd7
Amigo2
Papln
Dcn
Cacng5
Satb2
Wfs1
Pex5l
z-score (TPM)
C
AS AI PS PI
Ttr
Ighm
Prkcd
Zic1
Myoc
Tnnt1
Fst
Col6a1
Penk
Cartpt
Ntn5
PC2 (14.6%)
PC1 (32.3%)
D
Posterior vs Anterior
Ighm
Twist1
Trhr
Pou3f1
Fmod
Egr2
Myoc
Cartpt
Cpne7
Tnnt1
−log10(FDR)
log2FC
E
Cembrowski et al., 2016
This study
Dorsal
Ventral
Anterior
Posterior
Fst
Tnnt2
Egr2
Blnk
Vsig8
Pou3f1
Ighm
Gm10060
Ly6g6e
Rpl34−ps1
Ttr
Ntn5
Twist1
Trhr
Esyt3
Myoc
Fmod
Cpne7
Wnt4
Acta2
Spint1
Aldh1a2
Ngb
Otof
Nr2f2
Dio3
Upp1
Grp
Clec1a
Krt2
Stk32b
Pde1a
Sec1
z-score (TPM)
DG_d_3
DG_d_2
DG_d_1
DG_v_3
DG_v_2
DG_v_1
F
PS vs AS
Ighm
Ntn5
Nr2f2
Trhr
Twist1
Pou3f1
Cartpt
Ly6g6e
Gm13199
Cpne7
Vgll3
Tnnt1
G
PI vs AI
Tnnt1
Ighm
Ntn5
Shox2
Nr2f2
Pou3f1
Slc38a5
Egr2
Tnnt2
Trhr
Cartpt
H
Supra vs Infra
Fmod
Tnnt1
Myoc
Zic1
Cartpt
Aldh1a2
Egr2
Prkcd
Slc13a4
Ly6g6e

Figure S2

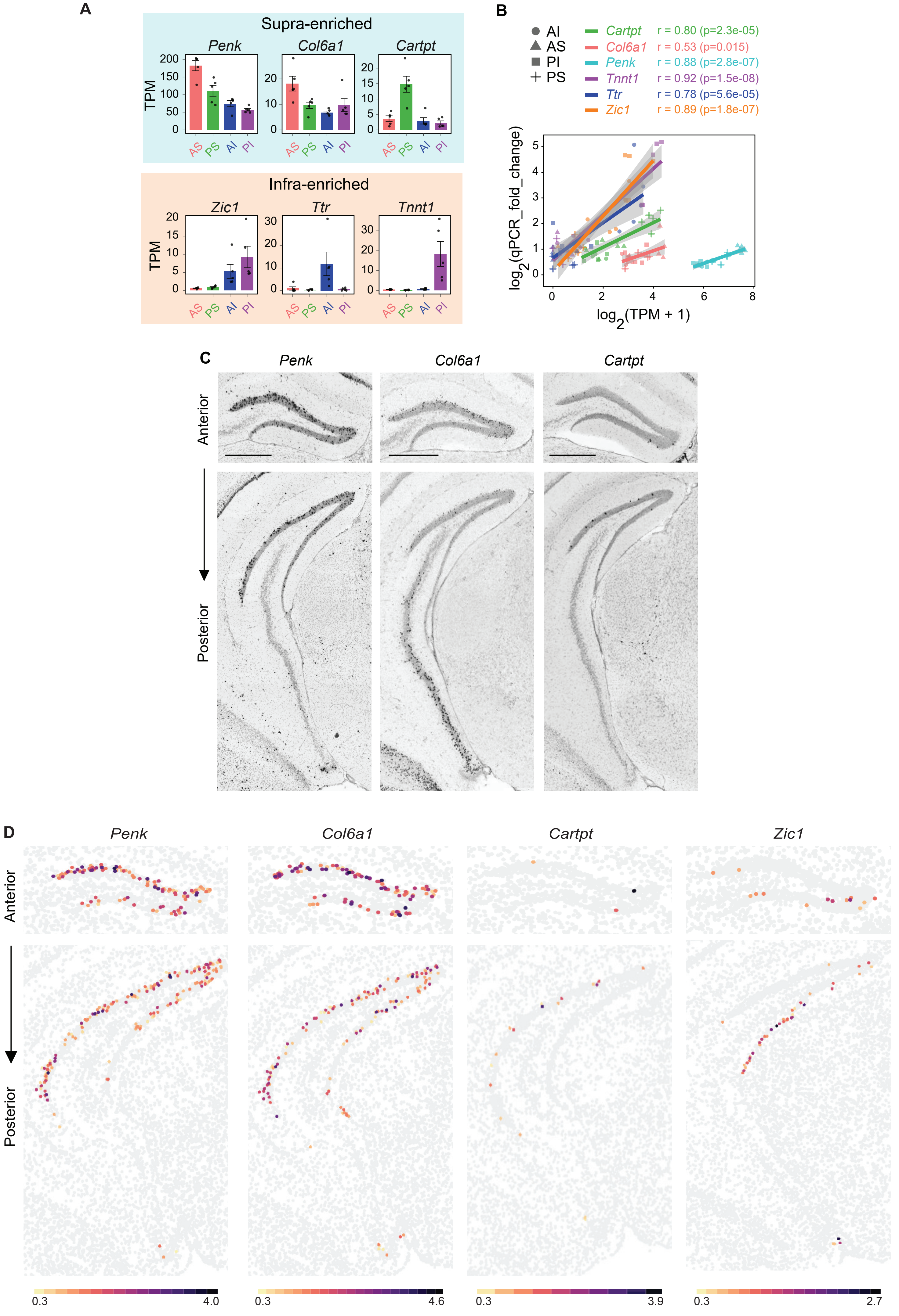
A
Supra-enriched
Penk
Col6a1
Cartpt
TPM
AS PS AI PI
Infra-enriched
Zic1
Ttr
Tnnt1
B
AI
AS
PI
PS
Cartpt r = 0.80 (p=2.3e-05)
Col6a1 r = 0.53 (p=0.015)
Penk r = 0.88 (p=2.8e-07)
Tnnt1 r = 0.92 (p=1.5e-08)
Ttr r = 0.78 (p=5.6e-05)
Zic1 r = 0.89 (p=1.8e-07)
log2(qPCR_fold_change)
log2(TPM + 1)
C
Penk
Col6a1
Cartpt
Anterior
Posterior
D
Penk
Col6a1
Cartpt
Zic1
Anterior
Posterior
0.3
4.0
0.3
4.6
0.3
3.9
0.3
2.7

Figure S3

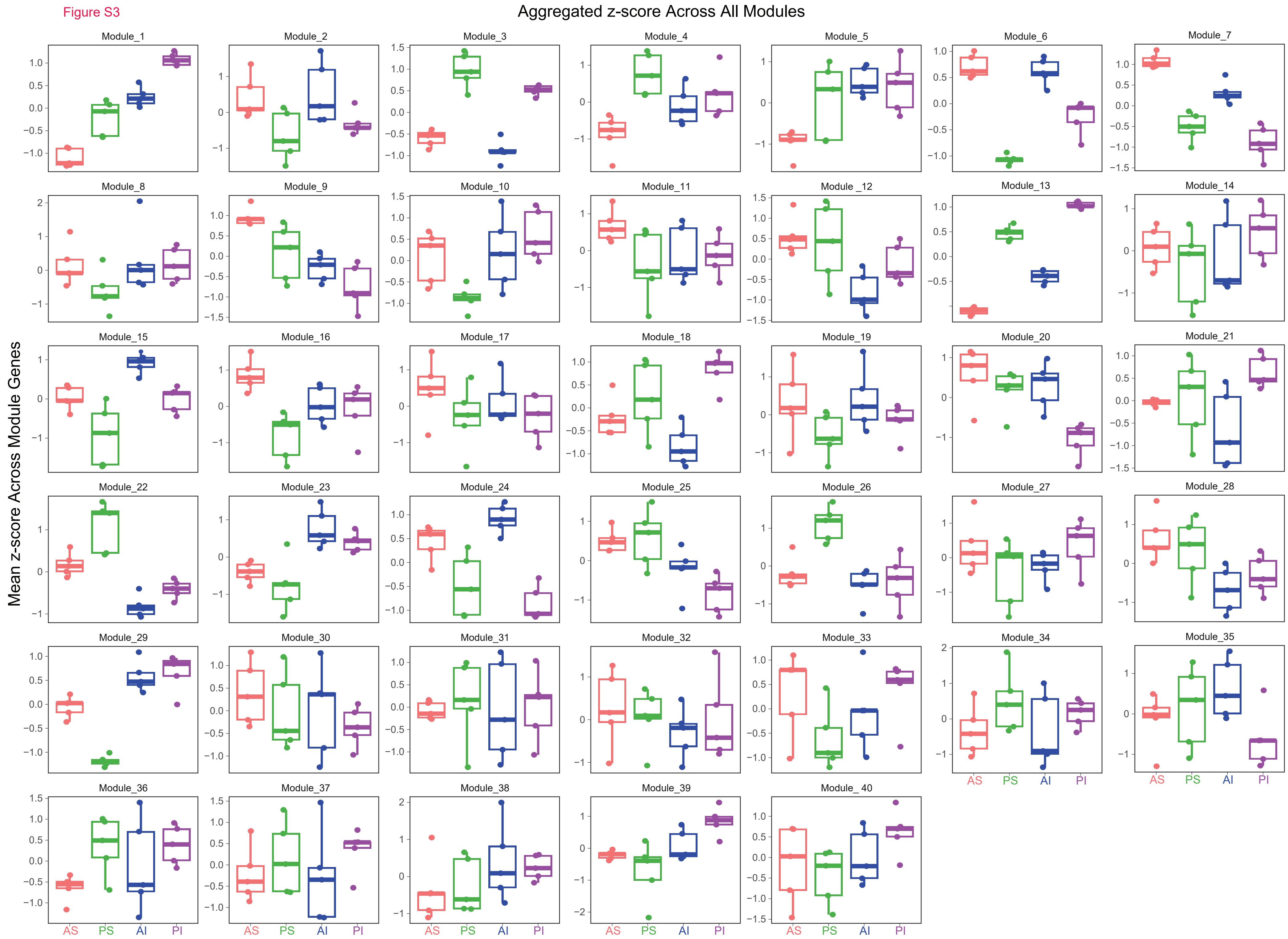
Aggregated z-score Across All Modules
Mean z-score Across Module Genes
Module_1
Module_2
Module_3
Module_4
Module_5
Module_6
Module_7
Module_8
Module_9
Module_10
Module_11
Module_12
Module_13
Module_14
Module_15
Module_16
Module_17
Module_18
Module_19
Module_20
Module_21
Module_22
Module_23
Module_24
Module_25
Module_26
Module_27
Module_28
Module_29
Module_30
Module_31
Module_32
Module_33
Module_34
Module_35
Module_36
Module_37
Module_38
Module_39
Module_40
AS
PS
AI
PI

Figure S4

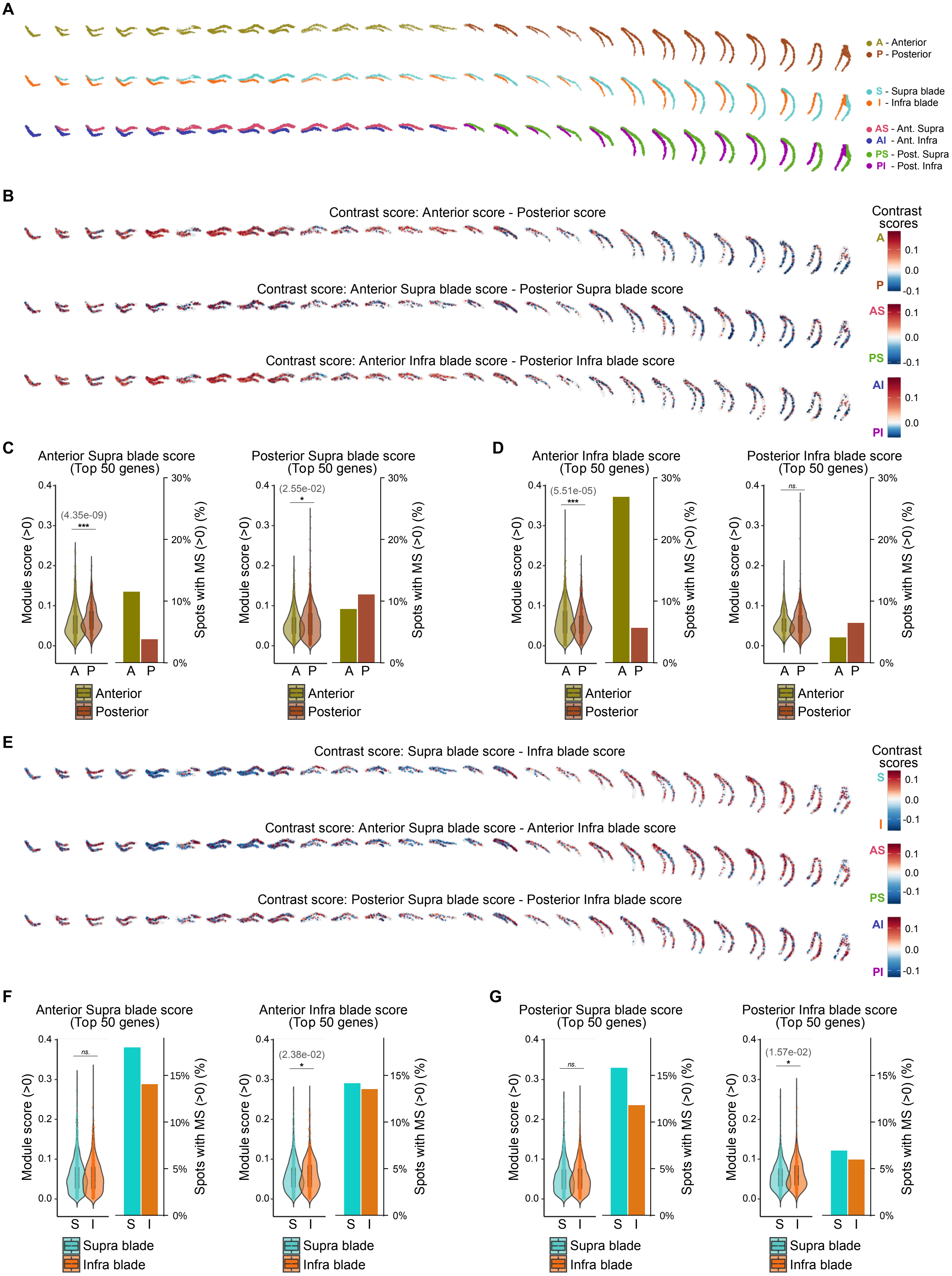

A
A - Anterior
P - Posterior
S - Supra blade
I - Infra blade
AS - Ant. Supra
AI - Ant. Infra
PS - Post. Supra
PI - Post. Infra
B
Contrast score: Anterior score - Posterior score
Contrast score: Anterior Supra blade score - Posterior Supra blade score
Contrast score: Anterior Infra blade score - Posterior Infra blade score
Contrast scores
C
Anterior Supra blade score (Top 50 genes)
(4.35e-09)
***
Posterior Supra blade score (Top 50 genes)
(2.55e-02)
*
D
Anterior Infra blade score (Top 50 genes)
(5.51e-05)
***
Posterior Infra blade score (Top 50 genes)
ns.
Module score (>0)
Spots with MS (>0) (%)
Anterior
Posterior
E
Contrast score: Supra blade score - Infra blade score
Contrast score: Anterior Supra blade score - Anterior Infra blade score
Contrast score: Posterior Supra blade score - Posterior Infra blade score
Contrast scores
F
Anterior Supra blade score (Top 50 genes)
ns.
Anterior Infra blade score (Top 50 genes)
(2.38e-02)
*
G
Posterior Supra blade score (Top 50 genes)
ns.
Posterior Infra blade score (Top 50 genes)
(1.57e-02)
*
Module score (>0)
Spots with MS (>0) (%)
Supra blade
Infra blade

Figure S5

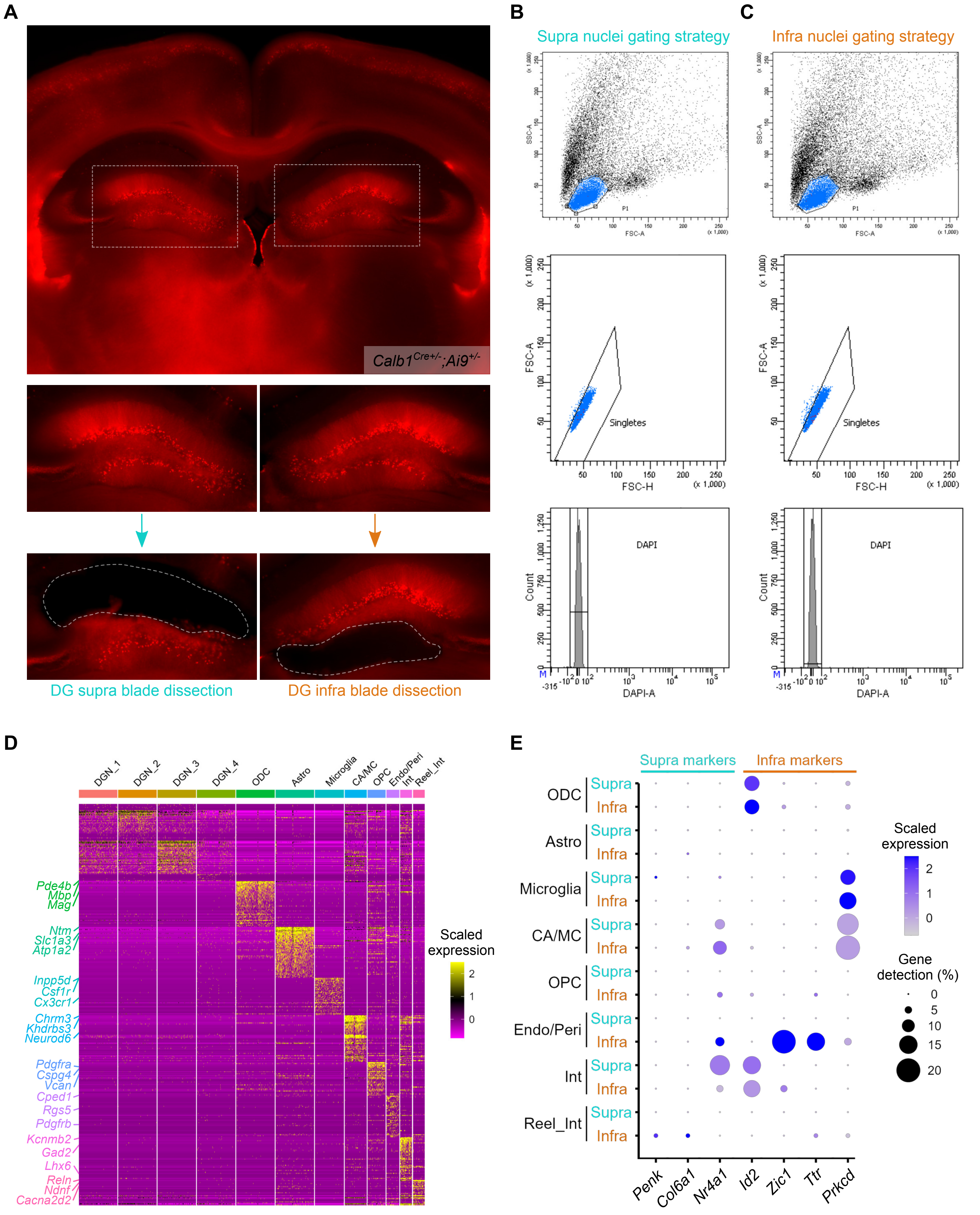
A
Calb1Cre+/-;Ai9+/-
DG supra blade dissection
DG infra blade dissection
B
Supra nuclei gating strategy
C
Infra nuclei gating strategy
SSC-A
FSC-A
P1
FSC-H
Singletes
Count
DAPI
DAPI-A
D
DGN_1
DGN_2
DGN_3
DGN_4
ODC
Astro
Microglia
CA/MC
OPC
Endo/Peri
Int
Reel_Int
Pde4b
Mbp
Mag
Ntm
Slc1a3
Atp1a2
Inpp5d
Csf1r
Cx3cr1
Chrm3
Khdrbs3
Neurod6
Pdgfra
Cspg4
Vcan
Cped1
Rgs5
Pdgfrb
Kcnmb2
Gad2
Lhx6
Reln
Ndnf
Cacna2d2
Scaled expression
E
Supra markers
Infra markers
ODC
Astro
Microglia
CA/MC
OPC
Endo/Peri
Int
Reel_Int
Supra
Infra
Penk
Col6a1
Nr4a1
Id2
Zic1
Ttr
Prkcd
Scaled expression
Gene detection (%)
